# Robust Group Fused Lasso for Multisample CNV Detection under Uncertainty

**DOI:** 10.1101/029769

**Authors:** Hossein Sharifi Noghabi, Majid Mohammadi

## Abstract

One of the most important needs in the post-genome era is providing the researchers with reliable and efficient computational tools to extract and analyze this huge amount of biological data, in which DNA copy number variation (CNV) is a vitally important one. Array-based comparative genomic hybridization (aCGH) is a common approach in order to detect CNVs. Most of methods for this purpose were proposed for one-dimensional profile. However, slightly this focus has moved from one- to multi-dimensional signals. In addition, since contamination of these profiles with noise is always an issue, it is highly important to have a robust method for analyzing multi-sample aCGH data. In this paper, we propose Robust Grouped Fused Lasso (RGFL) which utilizes the Robust Group Total Variations (RGTV). Instead of l_2,1_ norm, the *l_1_-l_2_* M-estimator is used which is more robust in dealing with non-Gaussian noise and high corruption. More importantly, Correntropy (Welsch M-estimator) is also applied for fitting error. Extensive experiments indicate that the proposed method outperforms the state-of-the art algorithms and techniques under a wide range of scenarios with diverse noises.

## 1. Introduction

Determination of changes in one- or multi-dimensional signals is of great importance and significance in fields such as Artificial Intelligence [1], Computer Networks [2] and Economy [3]. In addition to aforementioned areas, one- or multi-dimensional signals analysis has also found its way in biology and medicine [4]. Finding and specifying the place/time when a change is happening is significantly important especially when it comes to Copy-Number Variations (CNVs) in the genome [5]. The technique of microarray comparative genomic hybridization (arrayCGH or aCGH) paves the way for one to investigate genome-wide in order to capture all of these alternations [25]. Most of aCGH analysis methods were proposed for one-dimensional profile. However, slightly this focus has moved from one- to multi-dimensional signals where research are capable of capturing alternations from multi profiles simultaneously [7, 8]. Although multi-dimensional analysis brought more insight and information about aCGH landscape, most recently group approaches have been proposed for analyzing aCGH profiles [9]. There are two main advantages for group approach: 1) since these methods are considering a group of multi-dimensional profiles, they can provide us with more information about the structure of the profiles (where CNVs are occurring) and 2) this approach tends to be faster than previous methods as all profiles are considered at the same time [10]. Among all of the proposed methods for change point detection, Lasso [11] has found its way in numerous area from acoustics and speech processing [12] to compress sensing [13] and biomedical data analysis especially aCGH profiles [9]. Tibshirani et al. [14] proposed spatial smoothing and hot spot detection for CGH data using the fused lasso for one-dimensional aCGH analysis. Nowak et al. [5] proposed fused lasso latent feature model for multi-dimensional profiles.

Both of these methods applied the TV (Total Variation) norm for each sample independently and assume that the aCGH matrix is sparse. However, Zhou et al. [26, 25] proposed piecewise-constant and low-rank approximation (PLA) which assume this matrix is low-rank. Similar to previous methods, PLA is also take advantage of TV over for each sample. However, in order to deal with multi-dimensional features from a group perspective, Alvaiz et al. [10] proposed Group Fused Lasso (GFL) that utilizes Group Total Variation (GTV) for regularization. GTV uses the *l*_2,1_ norm instead of the *l*_1_. Bleakley et al. [9] utilized GFL with a weighted approach for multiple change point detection in aCGH profiles which lead to piecewise-constant approximations as well. While most methods are typically using first-order or alternating minimization schemes, Wytock et al. [16] proposed a fast Newton methods for the GFL that uses combination of a projected Newton method and a primal active set approach which is substantially fast.

### 1.1. Contribution

Based on *l*_1_-*l*_2_ and Welsch M-estimators, Robust Group Fused Lasso (RGFL) is proposed to recover true aCGH profile from noisy measurement. The proposed problem for RGFL is not convex, but its efficient solution is given by half quadratic (HQ) programming and its convergence is guaranteed. Due to the smoothness of the proposed algorithm, it is significantly faster than the other state-of-the-art algorithms which usually solve non-smooth problems. Extensive experimental results on simulated and real data illustrate the robustness of RGFL, especially when data are highly corrupted with non-Gaussian noise.

## 2. Robust Group Fused Lasso

### 2.1. Correntropy

To deal with non-Gaussian noise and impulsive noise, the concept of Correntropy is proposed in the realm of signal processing [17, 18]. Correntropy of two arbitrary variable *X* and *Y* is defined as

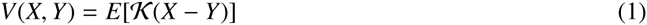
where *E*[.] is expectation operator and 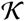 is a kernel function.

In practice, only finite number of samples 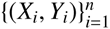 are available and the joint probability distribution of *X* and *Y* is usually unknown. Thus, Eq. (1) can be estimated as

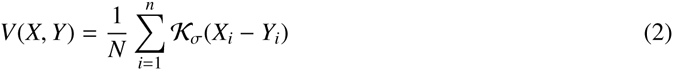
where 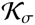 is Gaussian kernel function with the width of *σ* and 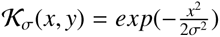. Correntropy is used to measure the similarity between *X* and *Y*. Liu et al. [17] induced a metric based on (2) for any two discrete vectors. They introduced the Correntropy induced metric (CIM) for any two vectors *x* ϵ *R^n^* and *y* ϵ *R^n^* as [18]

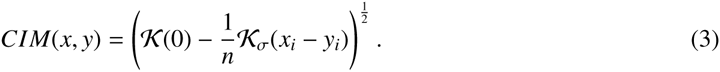

In contrary to mean square error (MSE), it should be noticed that Correntropy is a local similarity measure and the value of Correntropy is highly related to the width of Gaussian kernel function [17].

Correntropy can be applied in any noise environment because it has the probabilistic meaning of maximizing the error density function at the origin [17]. Further, if we choose Gaussian kernel function, Correntropy is a Welsch M-estimator in robust statistics [19].

### 2.2. Proposed formulation

Given a dataset *D ϵ R^n×m^* of aCGH profiles, it is desired to find the decomposition

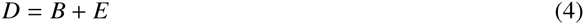
where *B ϵ R^n×m^* and *E ϵ R^n×m^* are true aCGH profiles and an additive arbitrary noise, respectively. A main problem in finding decomposition similar to (4) is that aCGH data are highly corrupted with various types of noise. Hence, the robust group fused lasso (RGFL) is proposed as follow

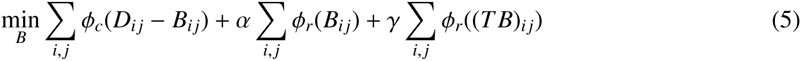
where *α, γ* > 0 are regularization parameters, 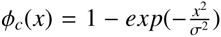 and 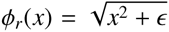 which is *l*_1_ *− l_2_* loss function in M-estimation and *T ϵ R^n−^*^1^*^×n^* is

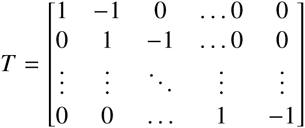
which makes *T B* takes the differences of the columns of *B*.

In fact, *ϕ_c_*(.) in Eq. (5) is a robust substitution of *l*_1_ − *l*_2_ in group fused lasso [20]. Although *l*_1_ is convex, but it is not differentiable at zero point and it results in oscillating around optimum and converging slowly toward the optimal solution. In contrast, *l*_1_ − *l*_2_ is differentiable arround zero point and can be solved significantly faster. Further, when *α* → 0 the *l*_1_ − *l*_2_ loss is equivalent to *l*_1_ loss. Hence, The second term in Eq. (5) forces the desired matrix *B* to be group sparse and the third term is a robust group total variation (RGTV). The first term is also CIM which is a robust measure for fitting error and can deal with diverse noise.

## 3. Solution of the proposed formulation

An iterative procedure to find the optimal solution of the proposed problem (5) is given by HQ minimization and the convergence of the solution is analyzed. Then, the optimal parameters for the proposed problem is given.

### 3.1. Half Quadratic Minimization

The proposed formulation (5) is not convex and cannot be solved directly. Fortunately, HQ minimization can be utilized to find the efficient optimal solution of Eq. (5) by an alternating procedure. Based on the conjugate function theory [21] and HQ theory [22, 23], the following lemma can be obtained.

#### Lemma 3.1.

*Let ϕ*(*x*) *is a loss function which satisfies the five conditions of additve HQ listed in [24]. Then, for any fixed x*

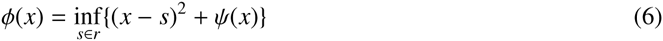

*where s is an auxiliary variable which is determined by the minimizer function δ*(.) *related only to ϕ*(.) *(see Table 1).*

**Table 1:**
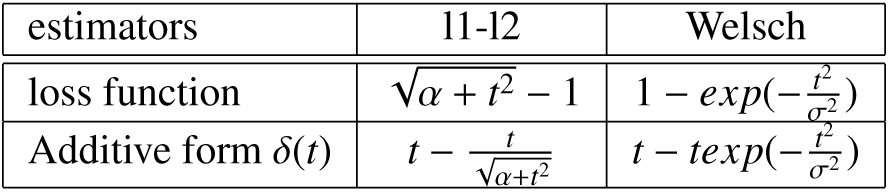
HQ loss functions and their corresponding minimizer functions

Replacing *phi*(.) in Eq. (5) by Eq. (6), we obtain the following augmented cost function

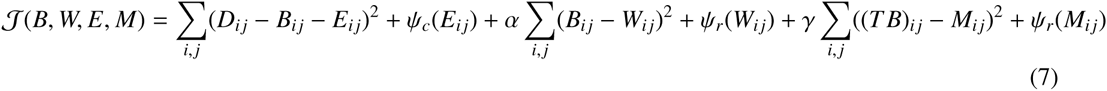
where E, *W* and *M* are HQ auxiliary variables. According to HQ, 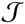 can be optimized by alternating minimization procedure. The HQ auxiliary variables are only related to the minimizer functions. Thus Σ *ψ*(*E_ij_*), Σ *ψ*(*W_ij_*) and Σ *ψ*(*M_ij_* become fixed and can be ignored for updating *B*:

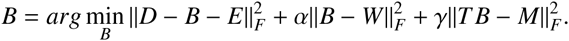

By derivating the above problem by *B*, we obtain

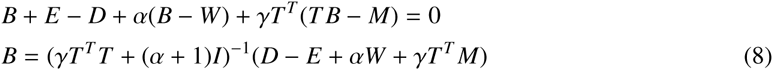
where *I* is the identity matrix. In short, the following steps must be taken for optimizing augmented cost function (7):

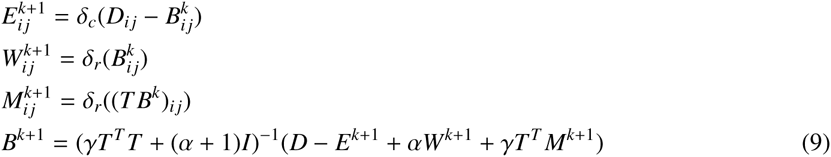

Algorithm 1 summarizes the procedure of RGFL.

#### Algorithm 1 Robust Group Fused Lasso

**Require:** D

1. *Initialize: D ϵ R^m×n^,* 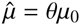*, B*_0_ *= B*_−1_ *=*0, *E*_0_ *= E*_−1_ *=* 0, *Z*_0_ *= Z*_−1_ *=* 0, *Y =* 0, *t*_0_ *= t*_−1_ *=* 1,
2. **while** not converge **do**
3. 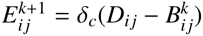
4. 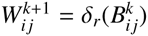
5. 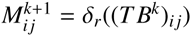
6. *γT^T^M^k^*^+1^) **end**
7. **return** *B*

### 3.2. Convergence Analysis

For fixed *E*, *M* and *W*, it is readily seen that

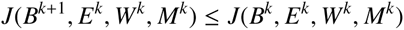

Further, according to the properties of HQ minimizer function *δ*(.) [24, 23] for a fixed *B*

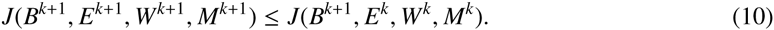

As Correntropy is bounded [17, 18], the sequence

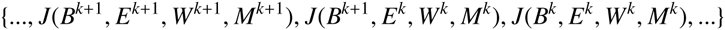
converges as *k* → ∞.

The inverse problem (8) can be solved using iterative method for linear equation system *Ax* = *b* that can even accelerate the convergence.

### 3.3. Parameter Selection

There are two parameters *α* and *γ* which can dramatically influence the performance of the proposed algorithm. In this section we go through a procedure by which these parameters are obtained. To do so, the data are divided into two subsets: *S*_1_ and *S*_2_. Let *Ρ*_Ω_(.) be the projection operator defined as

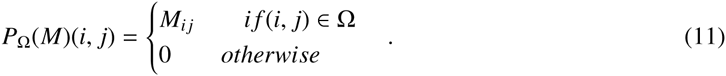

First, we perform the proposed method on the subset *S*_1_ with different values for *α* and *γ*, i.e.

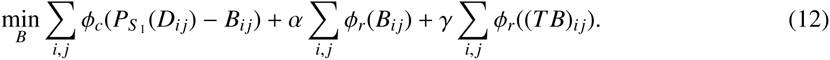

The solution for above problem is called 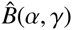 for each (*α, γ*) pair. Then, the pair is selected which minimize the predicion error defined as [25]

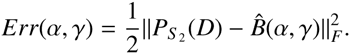

## 4. Experimental Results

In order to evaluate the proposed method, we performed our experiments on simulated and real data sets.

### 4.1. Simulated Data

In this section, the performance of the proposed method is compared with four well known methods including, TVSP [25], PLA [26], RCLR [8] and GFLseg [20]. In each case, 50 samples with 500 probes are generated according to the model (4) and various types of noise are added for investigation of the robustness. These noises are Uniform, Gamma, Beta and Laplace with settings tabulated in Table 2. Similar to [25], two types of aberrations are considered: the first type is added to each sample individually and the second type is shared among all samples at the same locations. The ratio between shared and total aberrations is denoted as *shared percentage*.

**Table 2:**
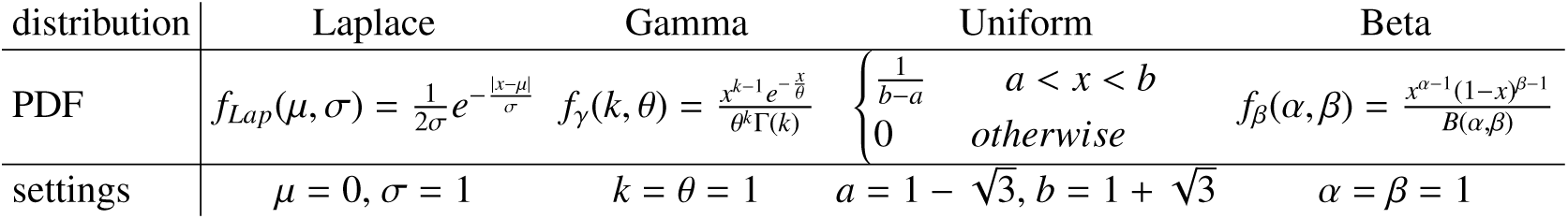
Probability density functions (PDF) and their paramters utilized as an additive noise to simulated data.

For each of these methods the Matlab implementations of them are downloaded and the parameters are tuned accordingly. Figure 1 illustrates the receiver operating characteristic (ROC) curves under Gamma, Uniform and Laplace noises and Figure 2 presents the comparison between stated methods in the case of accuracy under Beta noise.

**Figure 1:**
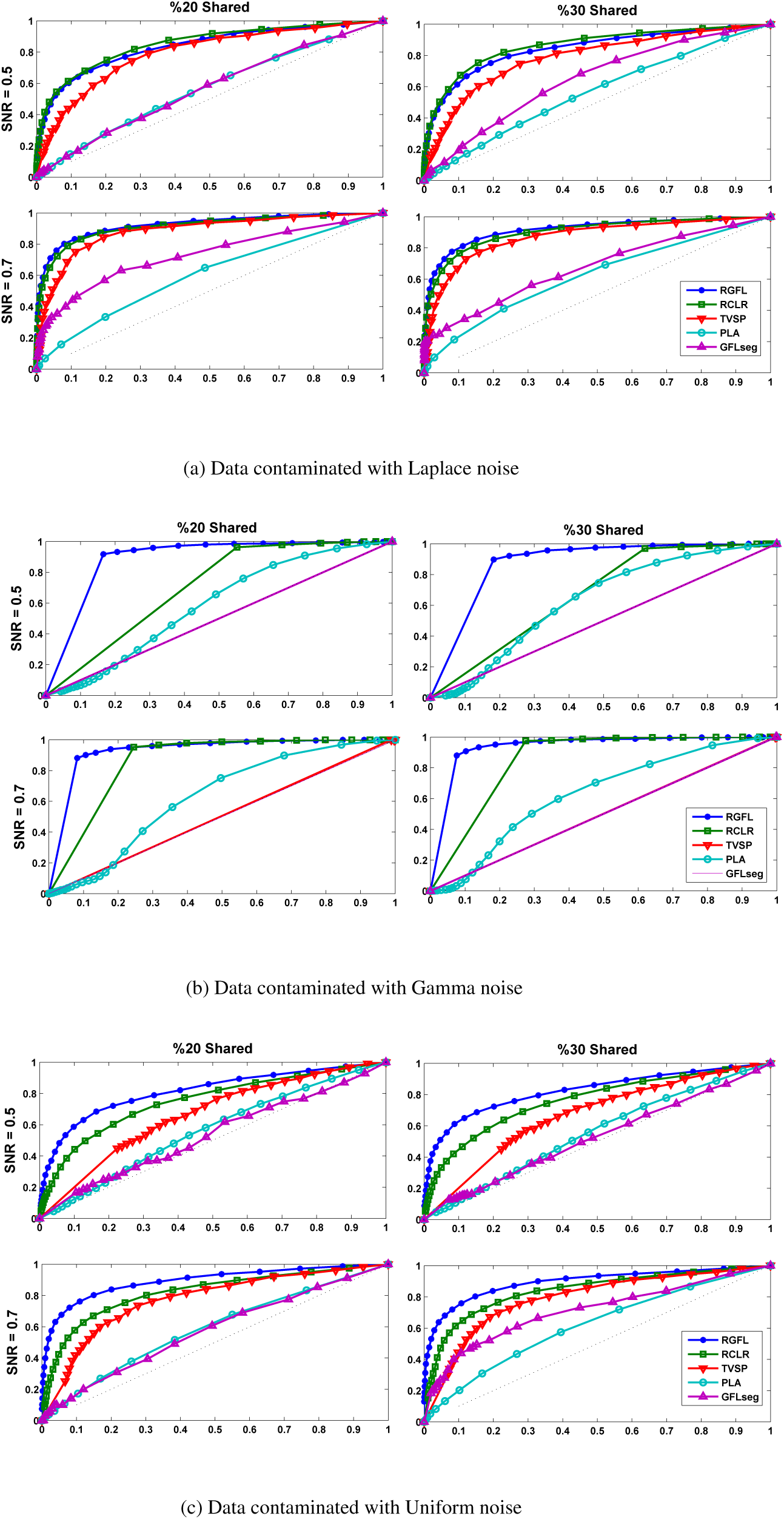
ROC for simulated data with non-Gaussian noise and different SNRs and shared percentages. Each row and each column are dedicated to a specific SNR and share percentage, respectively. The *x*-axis and *y* are false discorvery rate (FDR) and true discovery rate (TDR), respectively.++

**Figure 2:**
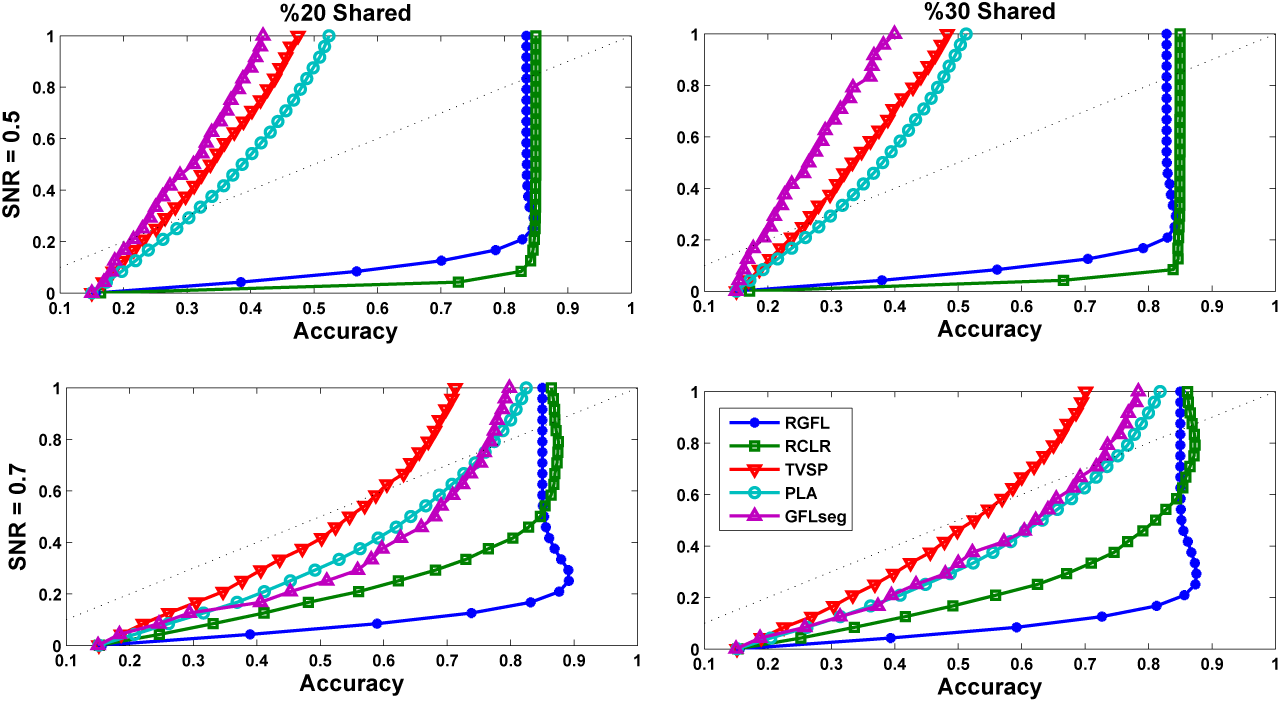
The comparison on simulated data from accuracy point of view. The data in this experiment are contaminated with Beta noise (see Table 2).

Obviously, more deviation from the diagonal line in ROC indicates better performance for the corresponding method. As plotted in this figure, the proposed method consistently and significantly outperforms PLA, GFLSeq and TVSP in all of the experimented noises. Especially for the higher SNRs and shared percentages which indicates that applying M-estimators and Correntropy makes the proposed method significantly robust against all of the studied noises and as a result much better performance in detection of common aberrations among multi-samples. As shown in Figure 1, RCLR seems to be extremely competitive, the reason for this observation is that similar to the proposed method, RCLR is also take advantage of Correntropy which makes it robust against uncertainty. However, despite this advantage of RCLR, the proposed method significantly and consistently outperforms it in Gamma and Uniform noises under both high and low SNR and shared percentage. This means utilizing Correntropy and M-estimators simultaneously, provides the proposed method with more robustness against uncertainty. In the case of Laplace noise with high value for SNR, RCLR appears to be slightly better than the proposed method, however, for lower SNR the proposed method again outperforms the RCLR.

These conclusions and observations are also deducible in Figure 2 for the accuracy under Beta noise. As illustrated in this figure, the proposed method significantly outperforms PLA, GFLSeq and TVSP in both high and low SNRs and shared percentages and gets extremely competitive with RCLR in higher SNR values. However, the proposed method achieves better accuracy for both high and low shared percentages when SNR is lower in comparison with RCLR.

### 4.2. Real Data

We investigate the performance of the proposed method on two independent real data sets about breast cancer. The Chin data set [27] includes profiles of 2,149 DNA clones for 141 primary breast tumors and the Pollack data set [28] has 6,691 human mapped genes for 44 advanced primary breast tumors. The results of these data sets are brought in Figure 3. The recovered profiles are plotted by heat map on the top, the recovered profile for a random sample in the middle and bar diagram, which presents the sum of number of gains over all samples for each probe with a threshold equals to one, at the bottom. In this figure, discovered regions with high amplifications are highly probable to be important for breast cancer, i.e., probes 67-70 and 39-42 are discovered for the Chin data set [27] and probes 178-184 are for the Pollack data set [28]. Interestingly, several identified functionally crucial regions for breast cancer, i.e., transcription regulation protein PPARBP, the receptor tyrosine kinase ERBB2 and the adaptor protein GRB7 are located within the discovered regions via the proposed method. Moreover, as illustrated in this figure *log*_2_-ratio of the denoised signals for both Chin and Pollack data sets are significantly smooth.

**Figure 3:**
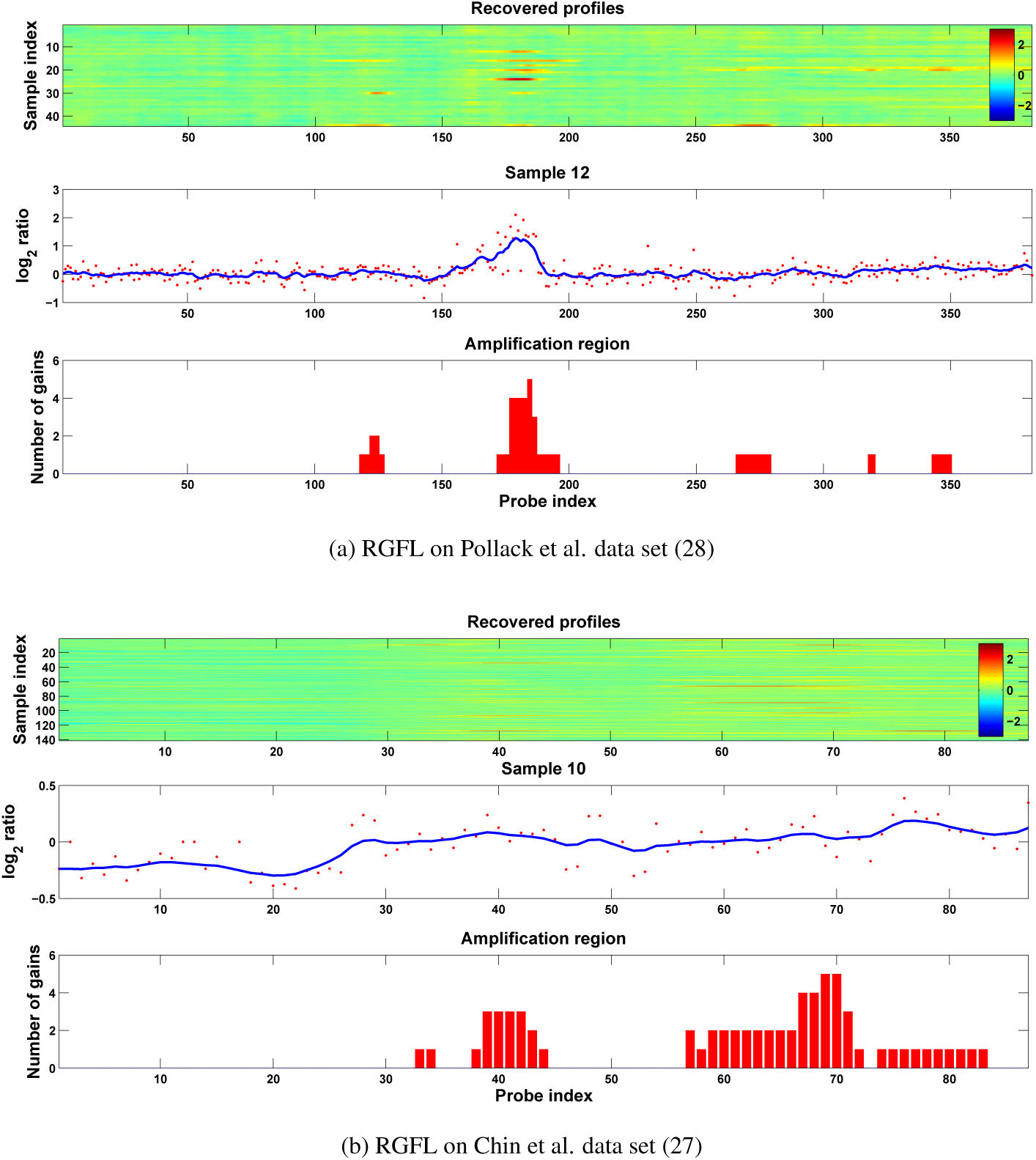
Heat and bar diagram for real datasets. (a) Performing RGFL on dataset introduced in [28] (b) Performing RGFL on dataset introduced in [27].

## 5. Conclusion

This paper presented Robust Group Fused Lasso (RGFL) to detect multiple changepoints in multi-dimensional signals. The proposed problem in this paper was non-convex, but an iterative procedure to find its efficient solution was given by HQ programming and its convergence analysis was given. In comparison to other state-of-the-art algorithms, the proposed method has shown significant robustness in dealing with high corruption, especially when data are mixed with non-Gaussian noise.

